# Automated Parameterization of Predictive Kinetic Metabolic Models from Sparse Datasets for Efficient Optimization of Many-Enzyme Heterologous Pathways

**DOI:** 10.1101/161372

**Authors:** Sean M. Halper, Iman Farasat, Howard M. Salis

**Affiliations:** Department of Chemical Engineering, Pennsylvania State University, University Park, PA 16802; Department of Biological Engineering, Pennsylvania State University, University Park, PA 16802

**Keywords:** pathway engineering, expression optimization, kinetic models, structured machine learning, biophysical models, design of experiments

## Abstract

Engineered heterologous metabolic pathways can convert low-cost feedstock into high-value products, though it remains a significant challenge to reliably and efficiently maximize end-product biosynthesis, particularly when many enzymes must be co-expressed together. When current approaches are applied to many-enzyme pathways, the construction and characterization process is highly iterative and laborious, while generating high-dimensional datasets that remain difficult to analyze for forward engineering efforts. To overcome these challenges, we developed a new algorithm that determines the highly non-linear and high-dimensional relationship between a pathway’s enzyme expression levels and its end-product productivity from common characterization of a small number of heterologous pathway variants. We combined kinetic metabolic modeling, elementary mode analysis, model reduction, de-dimensionalization, and genetic algorithm optimization into an automated procedure that parameterizes accurate kinetic metabolic models from sparsely characterized pathway variant libraries with varied enzyme expression levels. The resulting Pathway Maps are used to determine rate-limiting steps, predict optimal expression levels, identify allosteric interactions, rank-order enzyme kinetics, and prioritize protein engineering efforts. We demonstrate the Pathway Map Calculator algorithm on two experimental datasets, a 3-enzyme carotenoid biosynthesis pathway and a 9-enzyme limonene biosynthesis pathway, as well as a series of *in silico* pathway examples to rigorously demonstrate the algorithm’s accuracy, linear scaling, and high tolerance to measurement noise. By greatly reducing experimental efforts and providing quantitative forward engineering predictions, the Pathway Map Calculator has the potential to dramatically accelerate the engineering of many-enzyme heterologous metabolic pathways.

**Highlights:**

- We developed an automated algorithm that uses a small number of characterized pathway variants to determine the pathway’s expression-productivity relationship.
- The Pathway Map Calculator is accurate, scales linearly on many-enzyme pathways, distinguishes allosteric interactions, and tolerates substantial measurement noise.
- Pathway Maps are used to predict optimal enzyme expression levels, identify rate-limiting steps, and prioritize protein engineering efforts

## 1. Introduction

The metabolic engineering of organisms enables them to manufacture a wide variety of chemical products, but to achieve economic viability in a scaled-up process, the organism must have a sufficiently high yield, productivity, and titer. For commodities, specialty chemicals, and nutritional supplements, including 1,4-butanediol (Yim et al., 2011), isobutanol (Lan and Liao, 2011), succinic acid (Lee et al., 2006), n-butanol (Lim et al., 2013), lactic acid (Kong et al., 2015), isoprene (Zurbriggen et al., 2012), chondroitin (Wang et al., 2015), riboflavin (Lin et al., 2014) and cobalamin (Biedendieck et al., 2010), strict competition with existing processes requires that both the organism’s endogenous metabolic network and its heterologous pathways must be optimally tuned to maximize product synthesis, eliminate byproduct formation, and maintain robust organism growth rates in the expected bioreactor growth and media conditions. Higher-value products, such as erythromyocin (Jiang and Pfeifer, 2013), artemisinin (Ro et al., 2006), opioids (Thodey et al., 2014), taxadiene (Ajikumar et al., 2010), limonene (Alonso-Gutierrez et al., 2015), anthocyanins (Jones et al., 2017), and flavonoids (Trantas et al., 2009), often have more complex biosynthesis pathways, and therefore become more difficult to optimize because of the many enzymes needed to catalyze biosynthesis of the final product. In both cases, once a heterologous pathway has been introduced into an organism, and has been shown to minimally function, metabolic pathway optimization becomes an essential step to engineering an economically viable organism.

Heterologous metabolic pathway optimization is currently carried out by constructing and characterizing many pathway variants, incorporating different regulatory genetic parts and enzyme coding sequences to vary the enzymes’ expression levels and their intrinsic kinetics (Lynch et al., 2016; Oliver et al., 2013; Redding-Johanson et al., 2011; Smanski et al., 2014). Through combinatorial cloning and DNA assembly techniques, it has been possible to construct libraries of pathways utilizing different promoters, ribosome binding sites, and plasmid origins of replication to simultaneously vary the expression of multiple enzymes, modulating transcription rates, translation rates, and plasmid copy numbers (Scalcinati et al., 2012; Su et al., 2015; Watstein et al., 2015; Xu et al., 2013; Yu et al., 2016). Nowroozi et al. carried out pathway optimization on a nine-enzyme amorphadiene biosynthesis pathway in *E. coli*, characterizing 18 pathway variants with designed ribosome binding sites, resulting in a final titer of 3.6 g/L, including media optimization (Nowroozi et al., 2014). Zhao et al designed 81 pathway variants of the 3 enzyme (+)- catechin pathway, increasing transcription rates and using scaffolding proteins to increase titers by 128%, to a final titer of 910.9 mg/L (Zhao et al., 2015). Latimer et al used promoter libraries to construct and characterize 192 pathway variants of an 8-enzyme heterologous xylose utilization pathway in *Saccharomyces cerevisiae* for processing lignocellulose (Latimer and Dueber, 2017; Latimer et al., 2014).

More recently, it has become practical and common to integrate pathway modules into the host genome, and carry out pathway optimization directly on genomic genetic parts. Ng et. al. optimized a 5-enzyme Entner-Doudoroff pathway by combining the Operon Calculator and lambda red recombination to integrate rationally designed operons into the genome, followed by combining the RBS Library Calculator and MAGE genome mutagenesis to systematically vary enzyme expression levels, resulting in a 25-fold improvement in the NADPH regeneration rate (Ng et al., 2015). Li et al. utilized Cas9-assisted DNA repair to integrate a 14-enzyme β-carotene biosynthesis pathway into the *E. coli* genome, followed by construction and characterization of 103 pathway variants, 12 additional modifications to central metabolism, and media optimization to achieve a β-carotene titer of 2.0 g/L (Li et al., 2015). Through genome analysis and bioprospecting, libraries of homologous enzyme coding sequences have also been expressed and characterized with the goal of identifying enzymes with improved solubilities and kinetics in a destination host organism (Atsumi et al., 2010; Lanza et al., 2014; Shiue and Prather, 2014).

Collectively, these state-of-the-art examples present an interesting design challenge, which can be solved via algorithmic analysis and model-based prediction. The challenge is to identify the enzyme expression levels of a heterologous pathway that will maximize its productivity, given only a small set of characterized pathway variants with varied enzyme expression levels and measured end-product productivities. We focus on end-product measurements, both because of their commonality, and the difficulty of measuring intermediate intracellular metabolite concentrations and reaction fluxes. We also focus on pathway data-sets where enzyme expression levels were quantitatively varied with known changes, for example, by inserting libraries of promoters with well-characterized transcription rates or libraries of ribosome binding sites with well-characterized or well-predicted translation rates (**Figure 1**).

**Figure 1:**
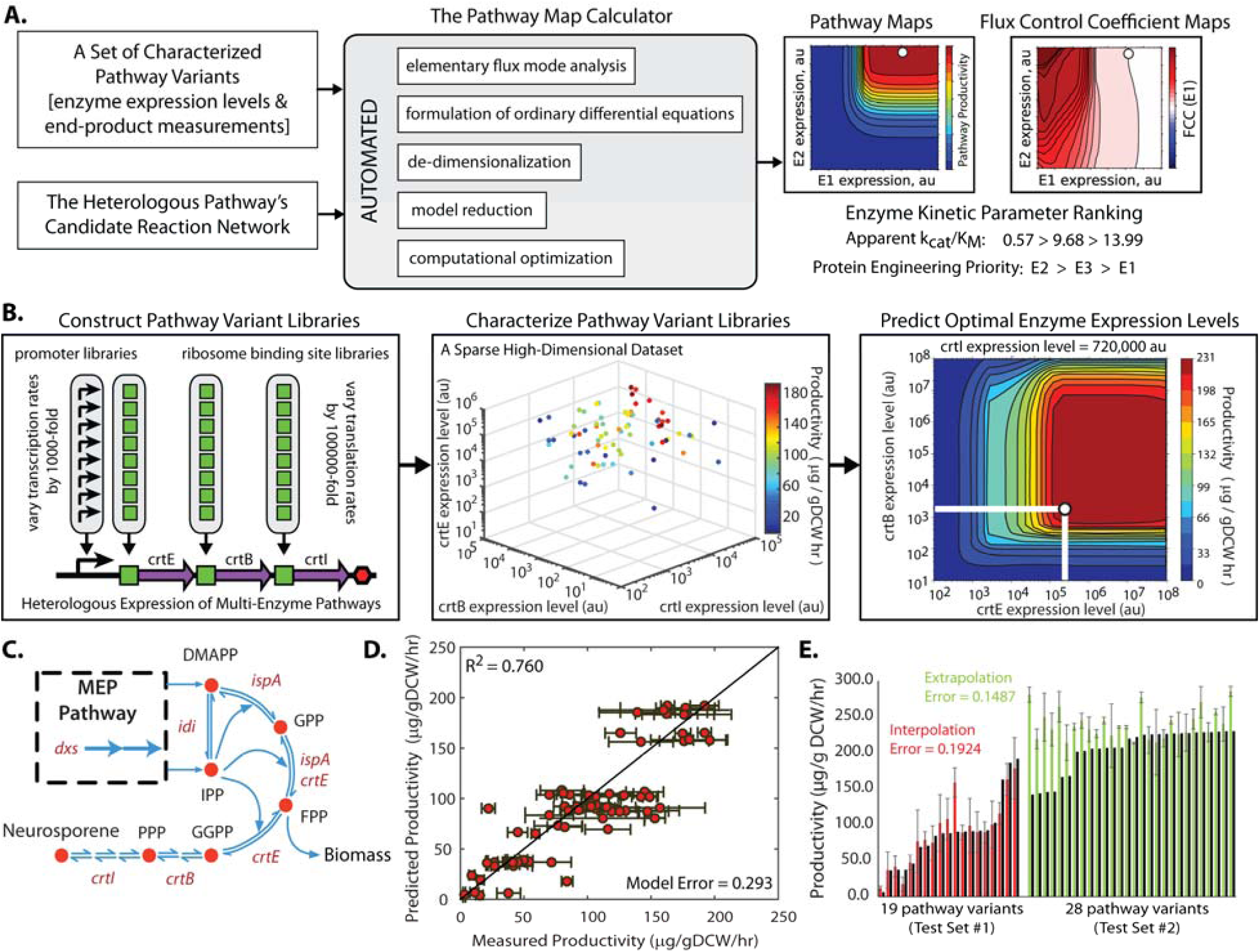
The Pathway Map Calculator workflow and its validation on a carotenoid biosynthesis pathway. (A) A dataset of characterized pathway variants and a candidate reaction network are fed into the Pathway Map Calculator. The algorithm uses the dataset to parameterize a kinetic metabolic model, which is then used to predict the pathway’s end-product productivity (a Pathway Map) and its rate-limiting steps (a Flux Control Coefficient Map) across all possible enzyme expression levels. (B) The workflow is illustrated on a 3-enzyme carotenoid (neurosporene) biosynthesis pathway, where 73 CrtEBI pathway variants were constructed and characterized, followed by using the Pathway Map Calculator to generate a Pathway Map and predict optimal enzyme expression levels. (C) A candidate reaction network for the 3-enzyme carotenoid biosynthesis pathway is shown. (D) The accuracy of the Pathway Map is initially tested by comparing predicted and measured neurosporene productivities across 73-pathway variant training set. (E) The accuracy of the Pathway Map is further tested by comparing predicted and measured neurosporene productivities across 19 characterized pathway variants with CrtEBI expression levels that overlapped with the training sets’ expression levels (interpolation test set #1) and across another 28 characterized pathway variants with CrtEBI expression levels that were higher than the training sets’ expression levels (extrapolation test set #2). Data points and error bars represent the means and standard deviations of 2 independent productivity measurements from Farasat et. al. (2014).

For several reasons, this challenge becomes particularly difficult when optimizing many-enzyme heterologous pathways, such as natural product pathways. First, the enzyme expression levels of a highly optimized pathway can be very high or very low, across a 100,000-fold range, depending on the enzymes’ catalytic efficiencies. Second, because enzymes work together synergistically, a pathway’s overall productivity is only improved when multiple enzyme expression levels are collectively tuned. Third, the relationship between a pathway’s enzyme expression levels and its productivity is multi-dimensional and highly non-linear; changes in enzyme expression will only improve the pathway’s overall productivity if the enzyme’s catalysis is rate-limiting. Fourth, if a combinatorial approach is used to optimize a pathway, then the number of experimental measurements will vastly exceed the throughput of most analytical techniques, such as mass spectrometry. Characterizing a tiny fraction of these pathways will yield a low chance of finding the pathway’s optimal enzyme expression levels. Fifth, and finally, once the optimal pathway variant has been constructed and characterized, further strain modifications, changes to the growth media or the expression of additional enzymes can imbalance the previously optimized pathway, requiring additional rounds of optimization.

Here, we describe and validate an algorithm, the Pathway Map Calculator, that determines the non-linear relationship between a pathway’s enzyme expression levels and its end-product productivities (**Figure 1A**). Using this relationship, called a Pathway Map, we predict the relative enzyme expression levels that maximize the pathway’s end-product productivity. All predictions are quantitative, experimentally actionable, and re-usable for a variety of applications. The algorithm requires two types of inputs: first, an experimental data-set consisting of end-product measurements and relative enzyme expression levels for each characterized pathway variant (**Figure 1B**); and second, a candidate network that lists the enzymes’ reactions and the corresponding metabolite stoichiometries (**Figure 1C**). Inputted end-product measurements may be assayed using any proportional measurement, such as LC-MS, GC-MS, enzyme-linked assays, or fluorescent biosensors. Inputted relative enzyme expression levels may be derived from measurements (e.g. proteomics) or model predictions of the genetic parts’ activities (e.g. predicted translation rates). Using these inputs, the automated *in silico* workflow combines kinetic metabolic modeling, elementary mode analysis, de-dimensionalization, model reduction, and genetic algorithm optimization to identify the enzymes’ kinetic parameters that best reproduces the inputted pathway variants’ measured productivities, and correspondingly, create a Pathway Map that shows the entire non-linear and high-dimensional relationship between the pathway’s enzyme expression levels and end-product productivity (**Figure 1B**). The Pathway Map enables efficient optimization of the pathway’s enzyme expression levels, provides insights into the pathway’s rate-limiting steps, and facilitates prioritization of protein engineering efforts to improve the slowest enzymes’ kinetics.

The development of the Pathway Map Calculator algorithm was inspired by previous kinetic metabolic modeling efforts (Matsuoka and Shimizu, 2015; Miskovic et al., 2015; Saa and Nielsen, 2016; Stanford et al., 2013) and by Ensemble Modeling in particular (Tan et al., 2011; Tran et al., 2008), though there are several notable distinctions. First, most kinetic metabolic models were developed to predict reaction fluxes through endogenous metabolic networks, constrained by reaction thermodynamics, isotopic metabolic flux measurements, and the outcomes of enzyme knock-out or over-expression experiments (Burgard et al., 2003; Khodayari and Maranas, 2016; Khodayari et al., 2014; Ranganathan et al., 2010). Here, our objective is to engineer and optimize heterologous metabolic pathways, utilizing the smallest possible set of characterized pathway variants with varied enzyme expression levels and end-product measurements. To our knowledge, an algorithm does not exist to solve this challenge. Second, while kinetic metabolic modeling and optimization have been recently combined to predict beneficial genetic interventions to endogenous metabolic networks, for example, using the k-OptForce algorithm (Chowdhury et al., 2014), the qualitative knock-out, knock-down, and knock-up predictions require interpretation to convert to experimentally actionable genetic part selections. Here, our developed algorithm predicts quantitative increases or decreases to specific enzyme expression levels to achieve desired productivity improvements, and these predictions are immediately convertible to the selection of a promoter or ribosome binding site sequence with a corresponding increase/decrease in transcription rate or translation rate (Alper et al., 2005; Mutalik et al., 2013; Salis et al., 2009).

## 2. Theory and Calculations

### 2.1 Overview of the Pathway Map Calculator Algorithm

The Pathway Map Calculator algorithm has three inputs: first, a candidate reaction network *RN_i,j,m,e,_* which establishes the list of substrates (index *i*), reactions (index *j*), reaction mechanisms (index *m*), and enzymes (index *e*), that participate in the metabolic network of interest; second, measured productivities *P*_*l*_ of a small library of pathway variants (index *l*); and third, the corresponding measured/predicted enzyme expression levels *E*_*e,l*_ for each pathway variant. Additional optional parameters include any measured/predicted substrate uptake rates *v*_*in,j,*_ and any measured/predicted internal fluxes *v*_*set,j.*_

In the first model construction phase, the algorithm automatically generates a system of de-dimensionalized, reduced differential equations that quantify the relationship between the metabolic network’s enzyme expression levels and its metabolic fluxes. Model reduction minimizes the number of unknown model degrees of freedom by incorporating elementary mode constraints and enzyme mole balance constraints and by grouping together co-dependent model parameters to create bounded, unitless reaction reversibility parameters. In the second model parameterization phase, genetic algorithm optimization is used to determine the unknown model parameters, using the kinetic model, productivity measurements, and enzyme expression levels to evaluate the optimization’s objective function. Finally, the parameterized kinetic model is analyzed, including identification of optimal enzyme expression levels, rank-ordering of enzymes according to their kinetics, and visualization of the multi-enzyme expression-productivity space.

### 2.2 Automated Calculation of the Elementary Modes

We use elementary mode analysis to enumerate the candidate reaction network’s elementary flux modes, and to convert all known metabolic fluxes into model constraints. Under steady-state conditions, the net fluxes through each reaction must satisfy the following mole balance:

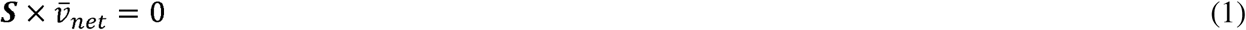

where the stoichiometric matrix ***S**_i,j_* is determined from the candidate reaction network and 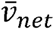 is the *j*×1 vector of net reaction fluxes. By calculating the null space of the stoichiometric matrix, the net reaction fluxes are then decomposed as a linearly independent set of reaction fluxes (Wagner, 2004),called elementary flux modes 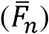, such that

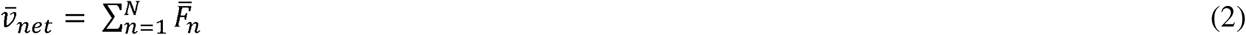

allowing us to equate the measured/predicted fluxes, such as the pathway productivities *P_l_*, to sets of individual reaction fluxes within the model. For example, a linear, unbranched pathway will have one elementary flux mode, and therefore requires only one productivity or substrate uptake measurement/prediction to specify the pathway’s net reaction flux. For more complicated metabolic networks, there will be a larger number (*N*) of elementary flux modes, requiring *N* flux measurements/predictions to fully specify all net reaction fluxes. Importantly, if there are insufficient flux measurements/predictions, then the remaining unspecified net reaction fluxes are considered degrees of freedom in the model parameterization. In cases where fluxes leaving the system, such as intermediate accumulation or competition with a different pathway are known, these fluxes should be included in the formulation of the reaction network, providing an elementary flux mode that accounts for the removal or build-up of metabolic flux at that point.

### 2.3 Automated Generation of the Kinetic Metabolic Model

We automatically generate the kinetic metabolic model, formulated as a system of ordinary differential equations, to quantify the rates of all reactions in terms of their metabolite and enzyme concentrations. First, each enzyme-catalyzed reaction is broken down into a set of reversible first-order and second-order reactions according to the enzyme’s reaction mechanism. We then quantify the reactions’ rates using elementary mass action kinetics according to the user-specified reaction mechanism. Over 30 reaction mechanisms are available for selection, including random-sequential, ordered-sequential, or ping-pong metabolite binding as well as several forms of allostery, including competitive, uncompetitive, and non-competitive inhibition.

For example, the reactions and fluxes for a mono-substrate, mono-product enzyme-catalyzed reaction are:

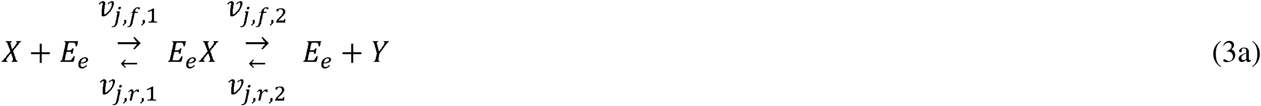

where *v*_*j,f*_ and *v*_*j,r*_ are the forward and reverse reaction fluxes for the j^th^ enzyme-catalyzed reaction using the e^th^ enzyme. The rates of these reactions are written as:

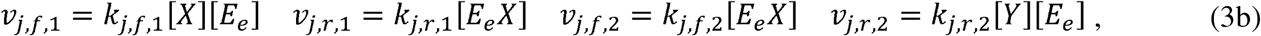

where *k*_1,*f*,1_, *k*_1,*r*,1_, *k*_1,*f*,2_, and *k*_1,*r*,2_ are kinetic rate constants, *[X]* is the concentration of substrate X, [Y] is the concentration of product Y, [*E*_*e*_] is the concentration of the e^th^ enzyme, and [*E*_*e*_*X*] is the concentration of enzyme-substrate complex. In another example, the elementary reactions and fluxes for the j^th^ bi-substrate, bi-product ordered-sequential reaction are:

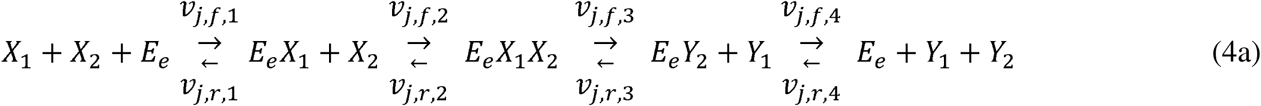

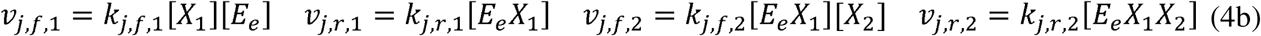

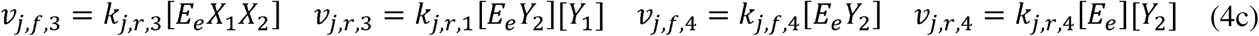

We then generate a system of ordinary differential equations for each metabolite, free enzyme, and bound enzyme complex in the reaction network, summing together all of the rate equations, yielding:

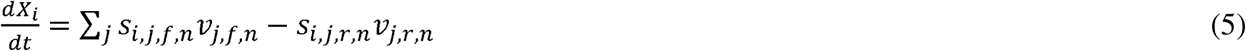

where the expanded stoichiometric matrices *s_i,j,f,n_* and *s_i,j,r,n_* describe the elementary reactions that produce or consume the i^th^ metabolite in the network, respectively, using the index *n* to count the number of elementary reactions to describe each reaction mechanism.

### 2.4 Automated De-dimensionalization of the Kinetic Metabolic Model

In the next step, we carry out automated de-dimensionalization to convert all metabolite and enzyme concentrations into dimensionless quantities. To do this, we select a reference pathway variant from the pathway library, which is any pathway variant that has particularly well-characterized end-product productivity measurement (smallest error bars) performed closest to steady-state conditions. We then multiply and divide all rate equations by the reference pathway’s corresponding metabolite and enzyme concentrations, following by regrouping and simplification. For the example of a mono-substrate, mono-product enzyme-catalyzed reaction, this procedure yields the following:

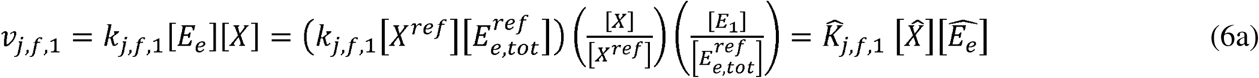

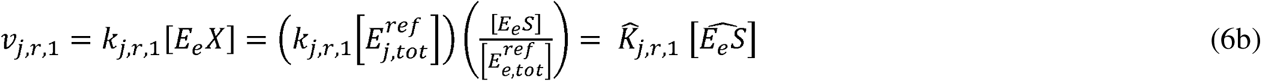

where [*X*^*ref*^] is the concentration of metabolite X in the reference pathway**,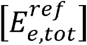** is the concentration of the e^th^ enzyme in the reference pathway, and **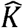** is the apparent kinetic rate parameter for each elementary step. All re-arrangements are carried out analytically. De-dimensionalized rate expressions are substituted into Equation (5).

Through de-dimensionalization, all metabolite and enzyme expression levels share the same relative scale, enabling us to (i) experimentally vary enzyme concentrations using several types of genetic modifications, e.g. changing promoters, ribosome binding sites, or plasmid copy numbers without algorithm modifications; and to (ii) compare relative changes in model-simulated metabolite productivities to relative changes in measured pathway productivities across different pathway variants. This approach was inspired by the ensemble modeling of kinetic metabolic models (Tan et al., 2011; Tran et al., 2008), which also performs de-dimensionalization of its metabolite and enzyme concentrations.

### 2.5 Automated Model Reduction using Enzyme Balances and Reaction Reversibilities

For a typical multi-enzyme pathway and candidate reaction network, the resulting system of de-dimensionalized differential equations will contain several unknown parameters. However, many of the parameters are co-dependent in that their values must collectively satisfy a set of mole balance and flux balance constraints. We therefore apply model reduction to eliminate all co-dependent parameters without altering the solution of the model equations. As a result, model reduction greatly reduces the number of pathway productivity measurements needed to identify the best-fit model, and accelerates model parameterization by placing sharp bounds on the parameter space. For example, for a 3-enzyme unbranched pathway with mono-substrate reaction mechanisms, there will be 22 unknown model parameters, including 12 kinetic rate constants, 9 metabolite/enzyme concentrations, and one net reaction flux. After model reduction, there will only be 10 unknown model parameters, including 6 bounded kinetic rate constants, 3 bounded metabolite/enzyme concentrations, and one net reaction flux.

First, we reduce the model by using the following set of enzyme mole balance constraints:

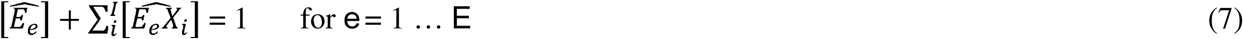

where the de-dimensionalized concentrations of the free enzyme and enzyme-substrate complexes will always sum to one. For a pathway with *E* enzymes, these constraints will reduce the number of unknown parameters by *E*.

Second, we reduce the model by using the following set of reaction flux balance constraints:

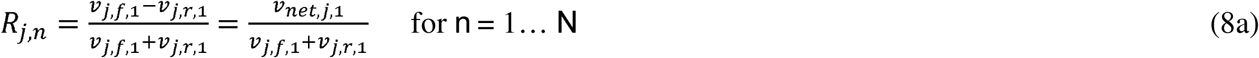

where *R*_*j,n*_ is the reaction reversibility of the n^th^ elementary reaction within the j^th^ enzyme-catalyzed reaction. A reaction reversibility is a dimensionless parameter that describes the net extent of a reversible reaction. If the reaction proceeds fully in the forward (reverse) direction, its reaction reversibility is one (negative one). Accordingly, we can re-formulate the forward and reverse reaction fluxes using a single reaction reversibility parameter, according to

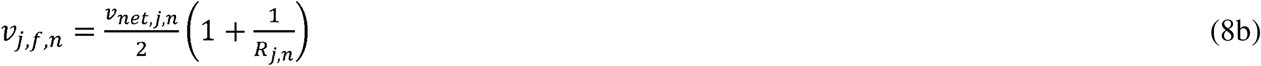

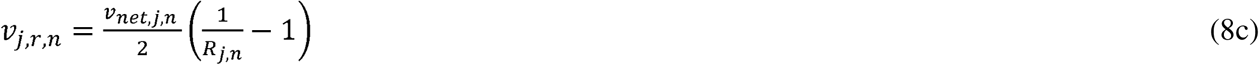

where all *R*_*j,n*_ are bounded by [-1,1]. Furthermore, if the thermodynamic favorability of a net reaction is known, then the values of the reaction reversibilities can be further constrained, using a variation of the flux force rule (Noor et al., 2014).

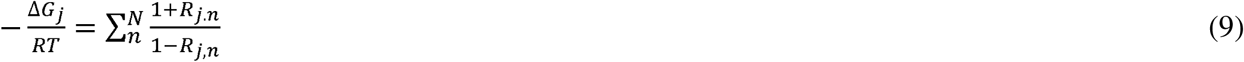

where Δ*Gj* is the free energy change for the j^th^ enzyme-catalyzed reaction, is the ideal gas constant, and is the temperature. When applied to the reference pathway, the relationships between the reaction reversibilities and the enzymes’ apparent kinetic parameters become:

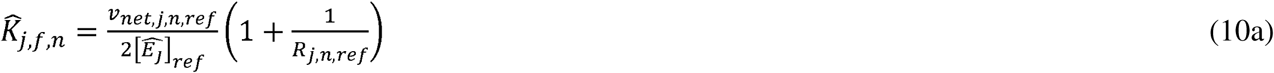

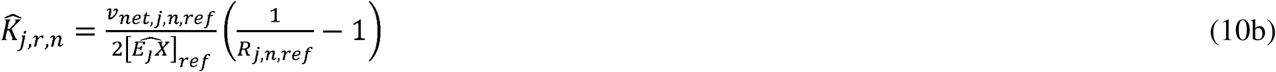

where these expressions become simplified because, for the reference pathway, **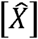** becomes one. By combining Equation 7 and Equation 10, we illustrate how the apparent kinetic parameters of a mono-substrate enzyme-catalyzed reaction will depend on the enzyme’s de-dimensionalized concentration and the two reaction reversibilities, according to:

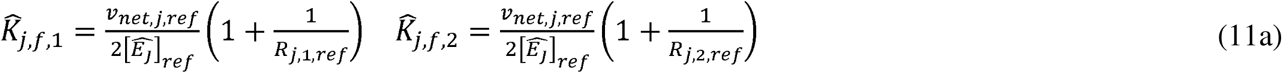

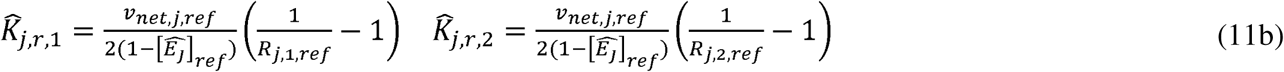

Therefore, the simplest *N*-enzyme pathway will require specification of only 3*N* unknown parameter values to determine the kinetic constants of the flux-constrained kinetic metabolic model. For more complex pathways, the number of unknown parameters will *at most* increase linearly with the number of enzymes.

### 2.6 Automated Parameterization of the Kinetic Metabolic Model

In the last step, we utilize the expression-productivity data-set to parameterize the kinetic metabolic model. Parameterization is composed as a many-variable scalar minimization problem, where the model’s degrees of freedom are the inputs and the objective function is the output to be minimized. The objective function *O* is an L_1_ norm that compares the pathways’ simulated net reaction fluxes and measured productivities, using the reference pathway’s corresponding quantities for normalization, according to:

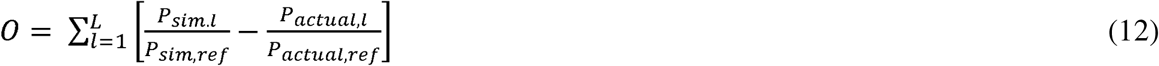

where *P*_*sim,l*_ is the simulated productivity of the *l*^th^ pathway variant and *P*_*actual,l*_ is the actual, measured productivity of the *l*^th^ pathway. Specifically, *P*_*sim,l*_ is the sum of net reaction fluxes that produce the product-of-interest, evaluated at the predicted/measured enzyme expression levels for the *l*^th^ pathway variant. By using simulations and measurements of the reference pathway to normalize these quantities, we may directly compare them on the same dimensionless scale.

We then accelerate the parameterization by carefully selecting the optimization algorithm and the model’s degrees of freedom. Individually, the characterized pathway variants will have different enzyme expression levels, and accordingly, different reaction reversibilities, reaction fluxes, and net fluxes through their elementary modes corresponding to the pathway variants’ measured productivities. Collectively, the enzymes in all pathway variants will exhibit the same apparent kinetic rate constants, 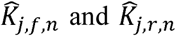. However, we do not select the kinetic rate constants as our model degrees of freedom because they are unbounded, forcing the optimization algorithm to search a large parameter space for a best-fit solution. Instead, as our model degrees of freedom, we select the reference pathway’s reaction reversibilities and de-dimensionalized enzyme concentrations, which greatly reduces and bounds the variable space. The reaction reversibilities are strictly bounded between -1 and 1, while the reference pathway’s enzyme concentrations vary between 0 and 1. We then use Equation 11 to inter-convert between these reference quantities and the enzymes’ apparent kinetic parameters.

We use genetic algorithm optimization to efficiently search this bounded space and identify sets of parameter values that best minimize the objective function in Equation 12. Our genetic algorithm implementation uses Gaussian-distributed mutation of variables and a uniformly random 2-point crossover for recombination, a mutation probability of 0.65, a cross-over probability of 0.35, a population size of 25 members. The algorithm is iterated at least 100 times. There are several aspects to this optimization problem where using a genetic algorithm will offer an advantage. The measured pathway productivities will always contain some level of experimental noise, which will convert a theoretically smooth objective function into a highly rugged space. With their ability to combine sub-optimal solutions via a recombination operator, genetic algorithms are particularly good at quickly converging to a global minimum even when many local minima exist. The rate of convergence and the quality of the fit can be readily improved by increasing the genetic algorithm’s population size and its number of iterations. The algorithm is also highly parallelizable with near 100% utilization of independent processors (cores) up to the size of the population. Moreover, we directly test the uniqueness of the parameterization process by utilizing a non-dominated selection operator, such as NSGA-II (Deb et al., 2002), to calculate the sets of model parameters that all equally minimize the objective function. Once optimization has been completed, we automatically confirm that the enzyme mole balance constraints and reaction flux balance constraints have been satisfied.

### 2.7 Automated Analysis of the Kinetic Metabolic Model

Once the best-fit kinetic metabolic model has been identified, we automatically carry out three types of analysis. First, we create a Pathway Map that quantifies the relationship between the pathway’s enzyme expression levels and its overall productivity. To do this, we carry out several *in silico* simulations using combinations enzyme expression levels that uniformly span the multi-dimensional expression space across a 100,000-fold range. We then use the reference pathway to convert the units of these simulated fluxes into measurable productivities, multiplying them by 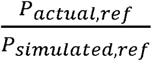. The Pathway Map shows how the pathway’s enzyme expression levels control the pathway’s productivity.

Second, we automatically create a FCC Map that quantifies the rate-limitingness of each enzyme and identifies the optimal enzyme expression levels that will yield maximum pathway productivity. Flux control coefficients (FCCs) are the first derivatives of pathway productivity with respect to each enzyme expression level (Fell, 1998). An enzyme is most rate-limiting when its FCC is one, and is not limiting when its FCC is zero. Importantly, the rate-limitingness of an enzyme is not intrinsic to the enzyme itself, but also depends on the other enzyme expression levels, according to:

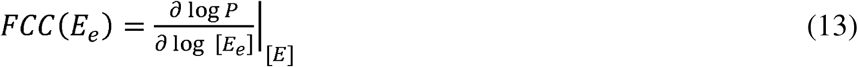

The global minimum of the FCC Map is the location where a pathway’s enzymes are least rate-limiting, and therefore where the pathway experiences its maximal productivity. We automatically designate these optimal enzyme expression levels, both on the Pathway Map and on the FCC Map. More comprehensively, the FCC Map can be used to identify the enzyme expression levels that yield *balanced* pathways and *optimally balanced* pathways. In a balanced pathway, all of the pathway’s enzymes are equally rate-limiting, carrying the same amount of reaction flux, and having the same FCCs. In contrast, an optimally balanced pathway is a special case where all of the enzymes have zero FCCs and where the only rate-limiting step is the precursor biosynthesis rate or substrate uptake rate. There are multiple combinations of enzyme expression levels that will yield balanced pathways, and with some types of allosteric regulation, multiple optimally balanced pathways can exist. It is also possible that implementing the optimally balanced pathway will exceed the host’s expression capacity, and therefore it becomes relevant to consider implementing the balanced pathways instead.

Third, we utilize the kinetic constants in the best-fit kinetic metabolic model to determine the enzymes’ apparent k_cat_ and K_M_ kinetic parameters in both out-of-pathway and in-pathway scenarios. These kinetic parameters are independent of the enzyme’s expression level, but do depend on its solubility and any acceleration of mass transfer, for example, via substrate channeling. To prioritize protein engineering efforts, we then automatically rank the pathway’s enzymes according to their in-pathway catalytic efficiencies (k_cat_/K_M_). To do this, we automatically generate a kinetic metabolic model that only contains the reactions catalyzed by the enzyme of interest, creating the corresponding system of differential equations. As before, these equations are de-dimensionalized in terms of substrate concentrations **,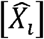** product concentrations **,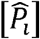**free and complexed enzyme concentrations, and lumped forward and reverse kinetic parameters, **,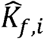** and **,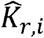**As an example, the following differential equations describe a mono-substrate, mono-product enzyme-catalyzed reversible reaction:

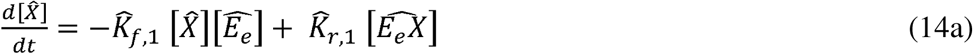

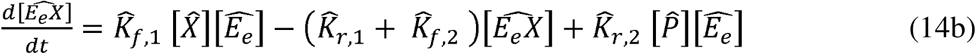

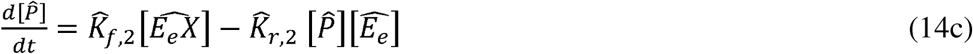

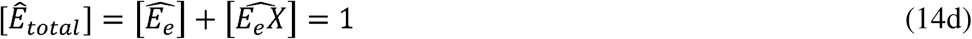

The rate law for the reaction is then automatically determined by first assuming a partial steady-state for the complexed enzyme concentrations, followed by substitution and simplification, yielding

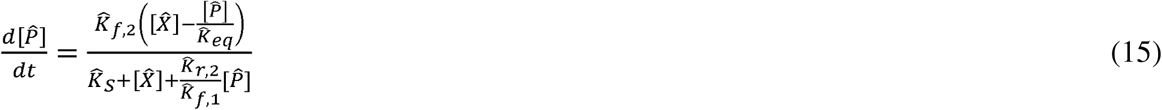

where **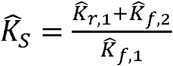** and **,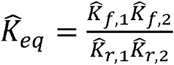** Here, we note that reaction is reversible and the rate of the reaction explicitly depends on the product’s concentration. If the product’s concentration is zero, the rate law is simplified, yielding an apparent **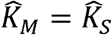** and **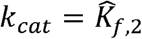**. However, if we consider the enzyme’s kinetics as they operate within the pathway, then the product’s concentration will not be zero. Instead, we define a useful scenario where the in-pathway’s substrate and product de-dimensionalized concentrations have reached a non-equilibrium steady-state, such that **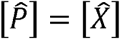**. In this scenario, simplification of the rate law yields an apparent 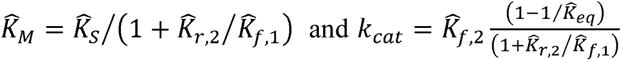. The same formalism is used to determine the apparent kinetic parameters for other types of reactions and reaction mechanisms under the two product concentration conditions (out-of-pathway and in-pathway scenarios).

## Results

## 3.1 Model Construction and Experimental Validation

The Pathway Map Calculator has been implemented in the Python programming language (v2.7) and uses the DEAP genetic algorithm package (Gagn, 2012) and the MPI for Python parallelization package (Dalcín et al., 2005). Unless otherwise noted, genetic algorithm runs were run with 25 members for 100 generations, and the results represent the best of three independent runs of the algorithm.

We first confirmed that the Pathway Map Calculator can generate predictive and accurate models of a pathway’s expression-productivity relationship, using a set of previously characterized 3-enzyme pathway variants from Farasat et al (2014). 73 pathway variants were constructed with rationally designed mutations to the ribosome binding sites controlling CrtEBI expression levels, employing the RBS Library Calculator to design the targeted mutations. The effects of these mutations on enzymes’ relative expression levels were then predicted using the RBS Calculator v2.0 (**Figure 1B**). Neurosporene titers and productivities were measured using absorbance assays and quantified with a calibration curve. We inputted the variants’ relative enzyme expression levels and neurosporene productivities into the Pathway Map Calculator, along with a candidate reaction network for neurosporene biosynthesis and biomass formation (**Figure 1C**). The resulting Pathway Map predicted the pathway’s 4-dimensional expression-productivity relationship, including the optimal enzyme expression levels that maximize the pathway’s neurosporene productivity. We then tested the Pathway Map’s accuracy by comparing predicted and measured pathway productivities, first for the training set of 73 characterized pathway variants, and then for a test set of 47 additional characterized pathway variants. The test set contains pathway variants that have unique combinations of expression levels either within the expression space of the training set (interpolation) or outside the training set’s sampled expression space (extrapolation).

First, we found that the Pathway Map’s predicted productivities were accurate to within 29.3%, on average, for the training set (**Figure 1D**), which was a lower fitting error over the previous modeling effort described in Farasat et. al. (2014). For the test set of pathway variants, we then found that the Pathway Map accurately predicted the pathways’ productivities to within 19.3%, on average, when predictions interpolated within the trained expression space (N = 19) (**Figure 1E**, red). Notably, the Pathway Map’s accuracy was retained or improved when productivity predictions extrapolated beyond the trained expression space with an overall accuracy of 14.9% (N = 28) (**Figure 1E**, green). The successful mapping of the CrtEBI pathway at a wide range of expression levels, both inside and outside the originally sampled expression space, confirms the Pathway Map Calculator’s ability to parameterize predictive kinetic metabolic models from a small number of characterized pathway variants.

As another example, we then applied the Pathway Map Calculator to a previously engineered 9-enzyme limonene biosynthesis pathway as developed by Alonso-Gutierrez et. al. (2015) where 23 pathway variants were constructed and characterized. 7 of the pathway’s enzyme expression levels were altered by inserting different promoters and vectors and by varying inducer concentration. In this example, the relative expression levels were directly measured using targeted proteomics, and the limonene titers and productivities were quantified using GC/MS according to a calibration curve. We then inputted the pathway variants’ measured enzyme expression levels (**Figure S1**) and limonene productivities (**Figure S2**) into the Pathway Map Calculator along with the candidate reaction network (**Figure 2A**). The resulting Pathway Map (**Figure 2B**) and FCC Map (**Figure 2C**) predict how changing the enzymes’ expression levels will affect the pathway’s overall limonene biosynthesis rate, including the optimal expression levels that are predicted to maximize limonene production (**Figure 2B**, white circles). Six informative two-dimensional plots are shown here; all 21 Pathway Map 2D plots and 42 FCC Map 2D plots are shown in **Figure S3**, **Figure S4**, and **Figure S5**.

**Figure 2:**
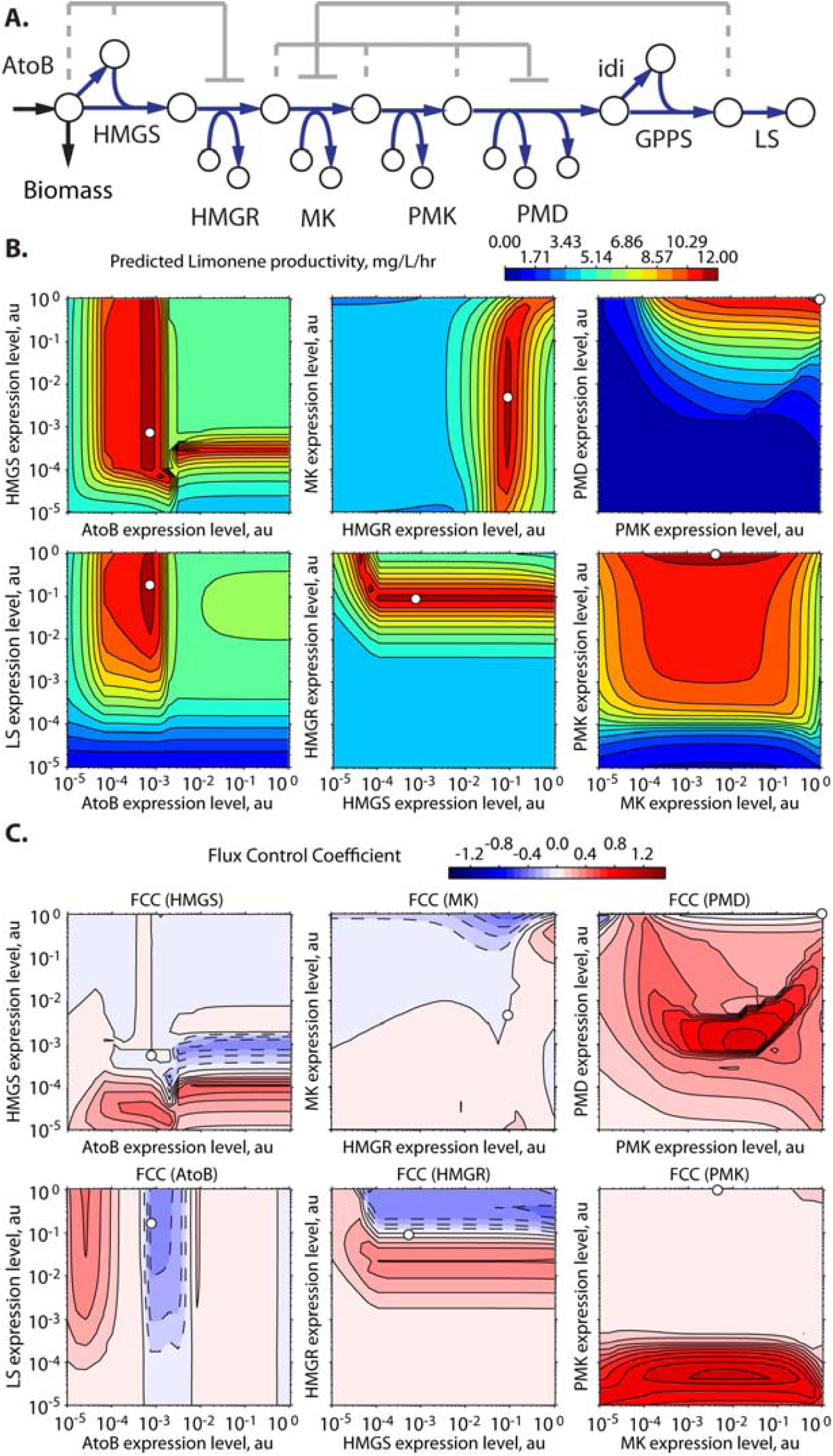
The Pathway Map Calculator is applied to an 9-enzyme limonene biosynthesis pathway with 23 characterized pathway variants as described in Alonso-Gutierrez et. al. (2015). (A) The candidate reaction network for limonene biosynthesis is shown, including three known allosteric interactions (dashed lines).(B) The generated Pathway Map predicts how changing enzyme expression levels will alter limonene productivity. The optimal expression levels are labeled (white circles). 2D slices of the 7D space are shown; the expression levels of the remaining 5 enzymes are kept constant at their optimal levels. idi and GGPS expression levels are fixed at measured values. (C) The generated Flux Control Coefficient Map shows how changing enzyme expression levels alters the rate-limitingness of the selected enzymes. A positive (negative) FCC indicates that expression of more (less) enzyme will lead to higher productivity. An enzyme’s optimal expression level will coincide with an FCC of zero. Allosteric interactions can lead to multiple local minima and maxima in expression-productivity space as indicated by multiple zero points in the FCC Map.

In agreement with Alonso-Gutierrez et. al. (2015), the Pathway Map predicts that limonene synthase (LS) is a key rate-limiting enzyme in the pathway, and that it should be expressed at a very high level. Specifically, the optimal LS expression level is 0.18 on the Pathway Map’s proportional scale, which varies from 10^-5^ to 1.0; increasing LS expression beyond this point is not predicted to increase limonene titer. Similarly, the Pathway Map predicts that the PMK and PMD expression levels should be maximally high (1.0 on the scale) because, besides increasing overall pathway flux, maximally increasing PMK and PMD expression will further reduce the concentrations of mevalonate-5P and mevalonate-5PP, which decrease MK-and PMD-catalyzed reactions via allosteric inhibition. However, because of the apparent differences in enzyme kinetics and the presence of allosteric inhibition, the optimal AtoB, HMGS, MK, and HMGR expression levels are predicted to be 235-fold, 235-fold, 38-fold, and 2-fold less than optimal LS expression level, respectively. Qualitatively, the analysis in Alonso-Gutierrez et. al. (2015) agreed that decreasing HMGS expression can lead to greater limonene biosynthesis.

Moreover, the Pathway Map provides a visual guide to understanding how small changes in enzyme expression level will affect the overall pathway’s flux. For example, varying MK expression around its optimal level is not predicted to significantly change limonene synthesis, whereas precise tuning of AtoB and HMGR expression is expected to be necessary to achieve maximal productivity. Enzyme expression sensitivities are quantified according to their flux control coefficients (**Methods**), which are visualized using the FCC Map (**Figure 2C**). A positive (negative) FCC indicates that expression of more (less) enzyme will lead to a higher pathway productivity, whereas a zero FCC indicates that the enzyme’s expression exists at a minima or maxima in expression-productivity space. Enzymes with large swaths of high (red) or low (blue) FCCs are key modulators of pathway productivity. Moreover, when product biosynthesis competes with organism growth, the FCC Map predicts how evolutionary processes are likely to break the pathway to improve growth rate; here, after the pathway has been optimized, lowering HMGR expression will have the most negative impact on limonene biosynthesis. Altogether, the Pathway Map and FCC Map provide actionable, readily implemented predictions to maximize limonene biosynthesis, for example, by inserting new promoters and ribosome binding sites designed to tune enzyme expression towards targeted optimal levels.

**Figure 3:**
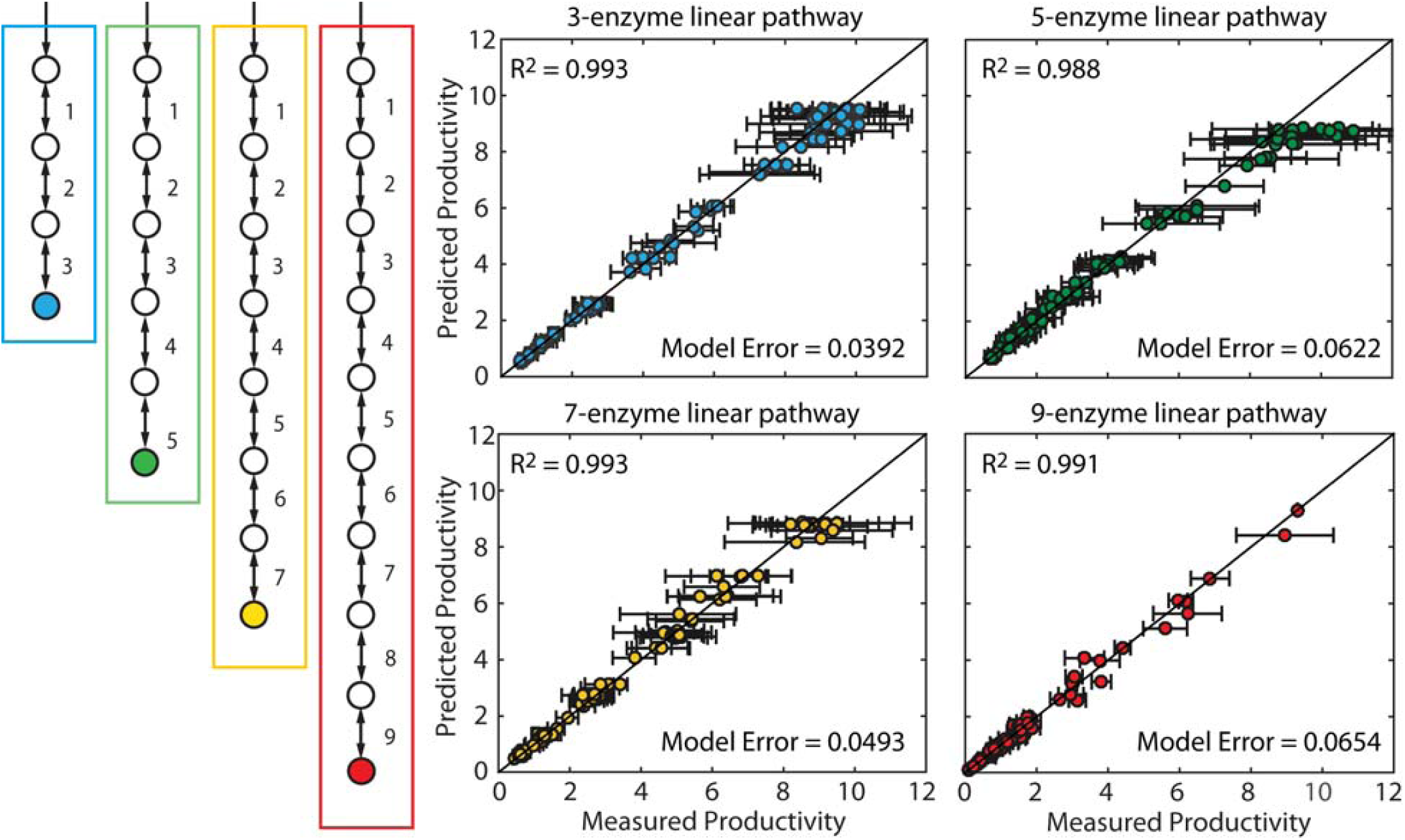
Pathway Map Accuracy versus Pathway Length. The productivities of four pathways with (blue) three, (green) five, (yellow) seven, and (red) nine enzymes were simulated with constant kinetic parameters, randomly assigned expression levels, and 10% simulated measurement noise. Productivities and expression levels from 100 *in silico* pathway variants were used to parameterize Pathway Maps, followed by comparisons between Pathway Map-predicted and *in silico*-generated productivities. Data points and error bars represent the means and standard deviations of three *in silico* simulations with varied enzyme expression levels and simulated experimental measurement noise.

### 3.2 Comparing Pathway Map Accuracy and Pathway Length

Industrially important pathways vary considerably in size and complexity, and therefore it becomes necessary to rigorously test the Pathway Map Calculator on many examples. To do this, the following sections describe rigorous accuracy and scaling tests carried out on a series of *in silico* simulated multi-enzyme pathways. To mirror the experimental construction and characterization procedure on realistic pathways, each pathway example has randomly generated enzyme kinetic parameters sampled from a physiological range. The end-product productivities of pathway variants are then simulated using randomly sampled enzyme expression levels across a 10,000-fold range, while keeping the enzyme kinetic parameters fixed. We also introduce simulated experimental measurement noise by multiplying the calculated productivities by a log-normally distributed random number with a median of one and a standard deviation of 0.10 (10% simulated measurement noise). These *in silico* pathway variants provide a realistic simulation of the sparse datasets acquired when experimentally characterizing libraries of pathway variants.

We first tested the predictive accuracy of the Pathway Map Calculator when faced with increasingly long unbranched pathways, from 3 to 9 enzymes, utilizing 100 *in silico* simulated pathway variants to parameterize Pathway Maps. For each of the progressively longer pathways, the algorithm was able to generate accurate Pathway Maps; compared to the training sets’ *in silico* calculated productivities, the Pathway Maps had fitting errors of 3.9%, 6.2%, 4.9%, and 6.5% for the 3, 5, 7, and 9-enzyme pathway, respectively. Small variations in fitting error are expected, due to the random generation of kinetic parameters for each pathway example, and the randomness of the enzyme expression level sampling. We then tested the accuracies of the Pathway Maps on interpolation and extrapolation test sets, and found that the prediction errors on the 3 to 9 enzyme pathways varied from 5.6% to 11.7% when predicting pathway productivities with expression levels that fell within the training sets’ space (**Figure S6**), and from 3.9% to 5.4% when predicting maximal productivities with optimal expression levels beyond the training sets’ space (**Figure S7**). Overall, we found that the Pathway Map Calculator was able to utilize only 100 characterized pathway variants to predict the optimal enzyme expression levels of unbranched pathways with at least 9 enzymes.

### 3.3 Comparing Pathway Map Accuracy and Pathway Variant Training Set Size

We next investigated the minimum number of characterized pathway variants needed to parameterize an accurate Pathway Map. We carried out simulations of the 9-enzyme unbranched pathway, generating *in silico* training sets with between 10 to 100 pathway variants. We then inputted these training sets into the Pathway Map Calculator, performing three independent algorithm runs, and comparing the Pathway Maps’ predicted productivities to the training set’s productivities (fitting error) as well as comparing the predicted productivities to a constant set of 125 pathway variants with *in silico*-generated productivities that broadly sampled the expression space (model error). We found that as the size of the training set was incrementally increased, the resulting Pathway Map’s overall model error dropped considerably with an apparent plateau when the training set size was 60 pathway variants (**Figure 4A**, green). Comparably, the fitting error stayed largely the same, regardless of the training set size, indicating that the Pathway Maps were not being overfit to the training set data (**Figure 4A**, red). As a point of reference for other pathways, the 9-enzyme pathway’s kinetic metabolic model has 27 unknown parameters; therefore, accurately parameterizing the model required about two times as many characterized pathway variants.

**Figure 4:**
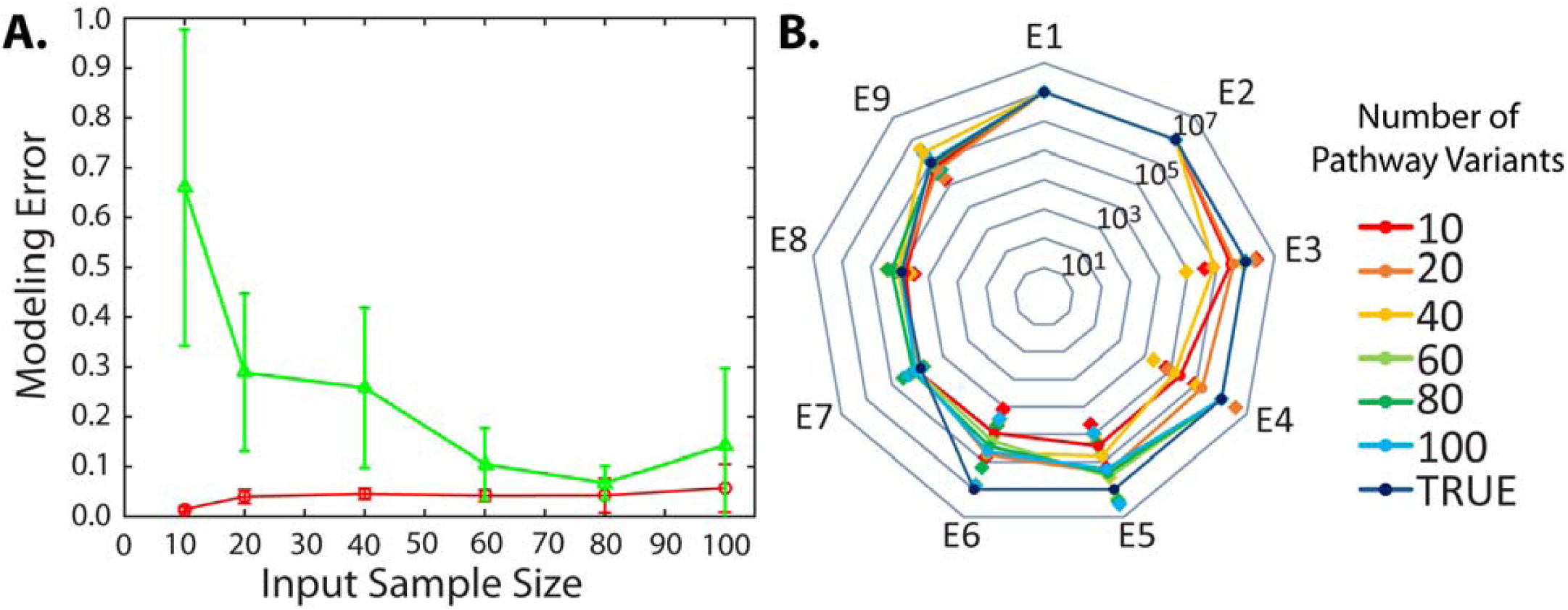
Pathway Map Accuracy versus Pathway Variant Training Set Size. (A) The (green) model error and (red) fitting error was determined when generating Pathway Maps of a 9-enzyme unbranched pathway with increasingly large pathway variant training sets. Error bars indicate the standard deviation of three independent algorithm solutions. (B) The corresponding Pathway Maps’ predicted optimal expression levels for each enzyme. Lines and circles represent the geometric mean of three independent runs of the algorithm. Diamonds represent the standard deviations. The true optimal enzyme expression levels are denoted.

We then assessed how the training set size affected the Pathway Maps’ predicted optimal enzyme expression levels, a key actionable prediction. The Pathway Map parameterized by the largest training set predicted optimal enzyme expression levels to within 2-fold of the true *in silico* determined values (**Figure 4B,** blue). As the size of the training set was reduced from 100 to 60, the predicted optimal enzyme expression levels did not appreciably change for seven of the enzymes (**Figure 4B**), though there was some variation in E5 and E6, due to the presence of a relatively flat plateau in the multi-dimensional expression-productivity space (**Figure S8**) that causes small changes in best-fit kinetics to have large changes in optimal E5 and E6 expression levels, though the predicted effect on pathway productivity was minimal. Overall, in this example, only 60 characterized pathway variants were needed to predict expression levels that achieve near-maximal pathway productivities.

### 3.4 Comparing Pathway Map Accuracy and Experimental Measurement Noise

We next investigated the Pathway Map Calculator’s ability to parameterize accurate Pathway Maps when trained on measurements with increasing amount of simulated experimental measurement noise. We carried out simulations of a 3-enzyme branched pathway with two end products, creating 104-pathway variant training sets. We then systematically varied the simulated multiplicative experimental measurement noise from 0 to 50% to generate different sparse datasets. We then inputted the training sets into the algorithm to parameterize Pathways Maps; the resulting differences in accuracy will reveal how measurement noise will affect Pathway Map accuracy (**Figure 5**). Without any simulated experimental measurement noise, the Pathway Map Calculator generated a Pathway Map with low model error (1.5% and 10% for products E and F, respectively). When the simulated measurement noise is typical of most analytical techniques (5 to 20%), the Pathway Map’s accuracy remained sufficiently low for actionable predictions (10% and 27% for products E and F at 20% simulated error, respectively). However, when the simulated measurement noise exceeded 20%, parameterization of the Pathway Map yielded noticeable skewed predictions with substantially higher model error (34.5% and 87.9% for products E and F at 50% simulated error, respectively). Overall, these comparisons show that the Pathway Map Calculator can generate accurate Pathway Maps, while accommodating the experimental measurement noise that is commonly found in most high-throughput analytical techniques.

**Figure 5:**
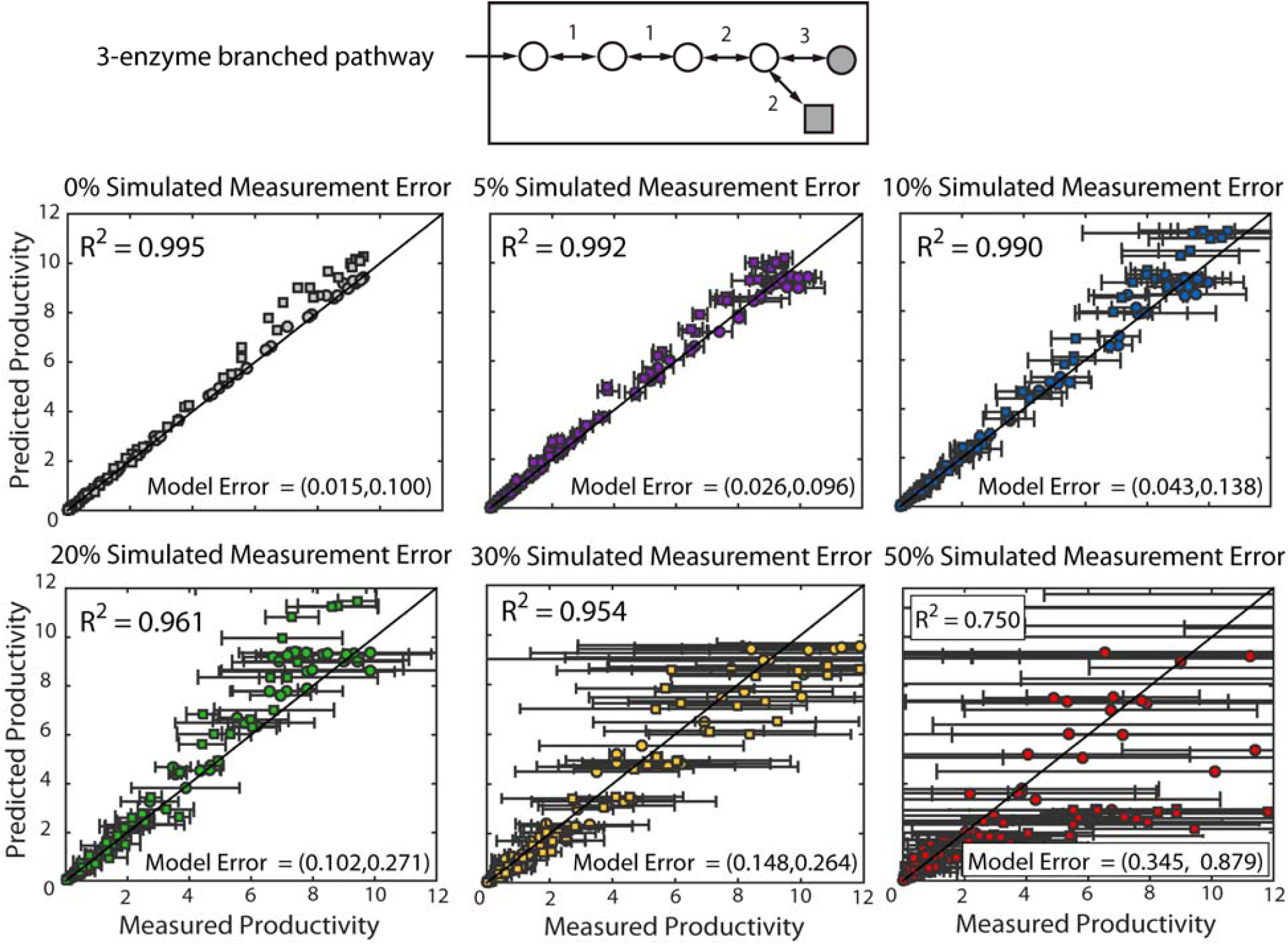
Pathway Map Accuracy and Experimental Measurement Noise. The productivities of a 3-enzyme, branched, two-product pathway were simulated using fixed kinetic parameters, randomly assigned expression levels, and log-normally distributed simulated measurement noise with standard deviations of (gray) 0%, (violet) 5%, (blue) 10%, (green) 20%, (yellow) 30% and (red) 50% of means. Pathway Maps were parameterized using training sets and their predicted productivities were compared to *in silico*-generated productivities. Data points and error bars represent the means and standard deviations of three independent algorithm runs. Plots with 30% and 50% simulated measurement noise show selected points for visual clarity.

### 3.5 Applying Pathway Mapping to Identify Allostery in Reaction Networks

The presence of enzyme allostery inside a pathway creates a non-linear, non-local form of regulatory control that often makes it challenging to engineer pathways for maximum productivity, particularly when it is not known when such allosteric interactions exist. Here, we show that the Pathway Map Calculator can use a small number of characterized pathway variants to identify the presence of allosteric interactions. To do this, we carried out simulations of a 3-enzyme pathway where the end-product inhibits the enzyme E2 via competitive inhibition (**Figure 6A**). The fixed training set consists of 100 *in silico* pathway variants with 10% simulated experimental measurement noise. We then created three candidate reaction networks – one without allostery, one with an incorrect form of allostery (feedforward regulation), and one with the correct form of allostery (end-product feedback inhibition) – and carried out independent runs of the Pathway Map Calculator using the fixed training set and the different candidate reaction networks (**Figure 6B**).

**Figure 6:**
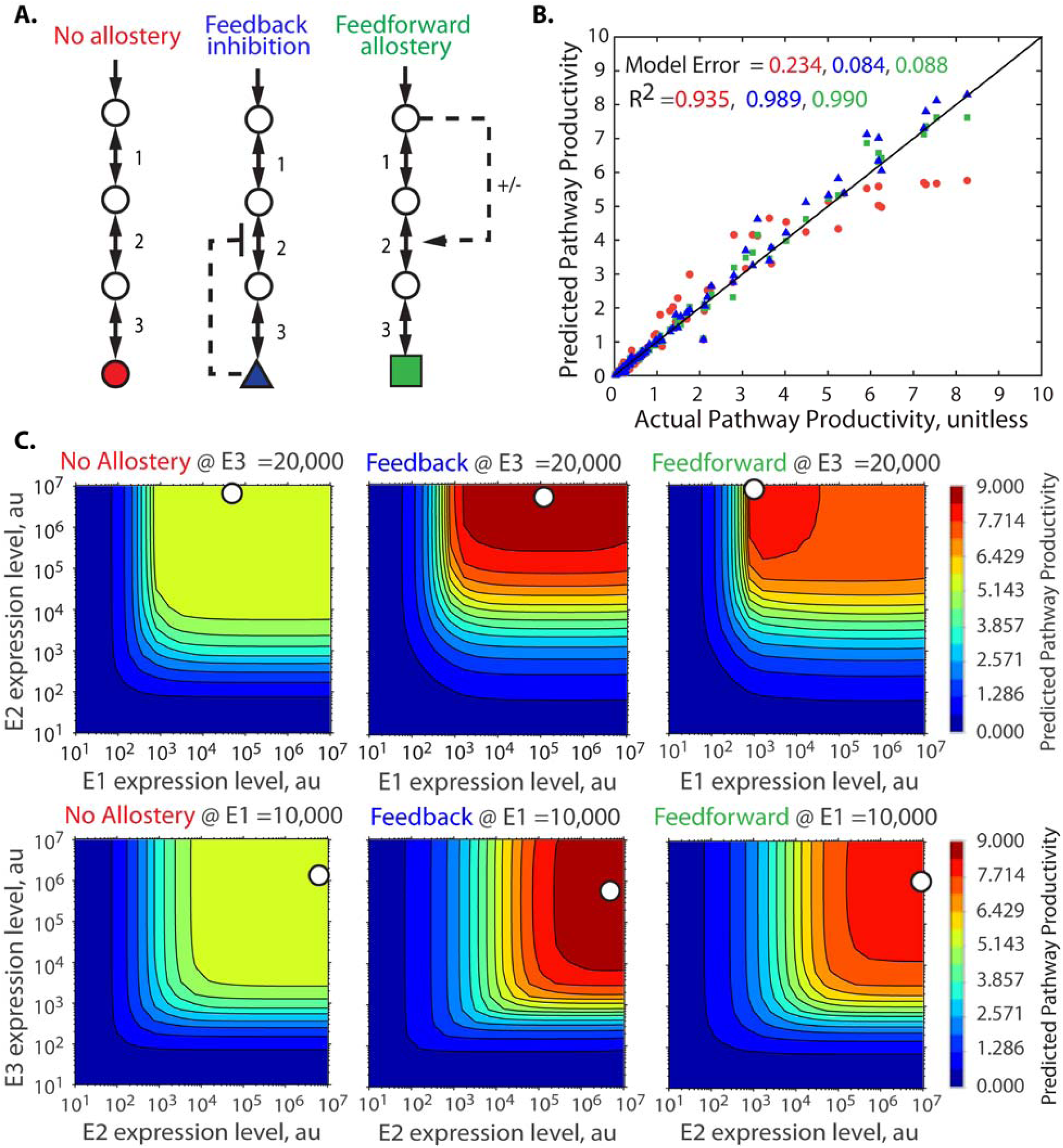
Identification of Allosteric Interactions using Pathway Mapping. (A) The same pathway variant training set is used with three different candidate reaction networks to parameterize Pathway Maps. (B) The Pathway Maps’ predicted pathway productivities are compared to the *in silico*-generated pathway productivities when using a candidate reaction network with either (red) no allostery, (blue) feedback allostery, or (green) feedforward allostery. (C) The parameterized Pathway Maps for the three candidate reaction networks, showing the differences in the expression-productivity relationship when different forms of allostery are assumed.

When the no-allostery candidate reaction network was used, the resulting Pathway Map has a high model error (23.4%) (**Figure 6C**). Without allostery, the pathway’s expression-productivity relationship has a large plateau, and the Pathway Map Calculator correctly determines that the training dataset can not be reconciled with the incorrect no-allostery candidate reaction network. However, when candidate reaction networks with allostery were used, the resulting Pathway Maps were accurate (model errors of 8.4% for feedback inhibition and 8.8% for feedforward regulation). Here, these different forms of allostery created very similar Pathway Maps (**Figure 6C**), and therefore the Pathway Map Calculator was only able to identify the correct candidate reaction network (with feedback inhibition) by creating a Pathway Map with a marginally lower amount of model error. Overall, by using model error as a proxy for the correctness of the candidate reaction network, the Pathway Map Calculator can distinguish between correct and incorrect candidate reaction networks, for example, with and without allostery. Importantly, when correct and incorrect candidate reaction networks both generate similar Pathway Maps, the Pathway Map Calculator can not readily distinguish them. However, in this scenario, the actionable predictions (optimal enzyme expression levels) to improve the pathway’s productivity are also similar.

### 3.6 Applying Pathway Mapping to Prioritize Protein Engineering Efforts

Protein engineering is frequently used to improve an enzyme’s intrinsic properties, such as solubility, turnover number (k_cat_), substrate selectivity, affinity (K_M_), and allostery, though the process of mutagenizing proteins and screening them for improvements can require a significant amount of time and resources. It therefore becomes important to first carry out protein engineering on the slowest enzymes, catalyzing the most rate-limiting steps in the pathway. Here, we show that Pathway Mapping identifies and ranks the apparent kinetic parameter values of a pathway’s enzymes, which can then be used to prioritize protein engineering efforts. We demonstrated this capability by carrying out simulations of a 3-enzyme pathway where the reaction reversibilities for E1 and E2 were about 1000-fold higher than those of E3. We then utilized a 100-pathway variant training set to parameterize a Pathway Map, followed by calculation of the pathway’s flux control coefficient (FCC) map, the pathway’s optimal expression levels, and the enzymes’ k_cat_ and K_M_ parameters. We also investigate whether an enzyme’s *apparent* kinetic parameters can change when the enzyme catalyzes a reaction in isolation (out-of-pathway) or together with the other enzymes in the pathway (in-pathway).

In this illustrative demonstration, we found that the pathway’s FCC map provides an ideal birds-eye view of the rate-limitingness of the pathway’s enzymes, showing the relationship between the enzymes’ expression levels and their flux control coefficients (**Figure 7A**). As before, we use the Pathway Map and FCC Map to predict the pathway’s optimal enzyme expression levels and we found that the enzyme E3 required the highest expression level to maximize pathway productivity, immediately suggesting that E3 should be prioritized for protein engineering efforts (**Figure 7B**).

**Figure 7:**
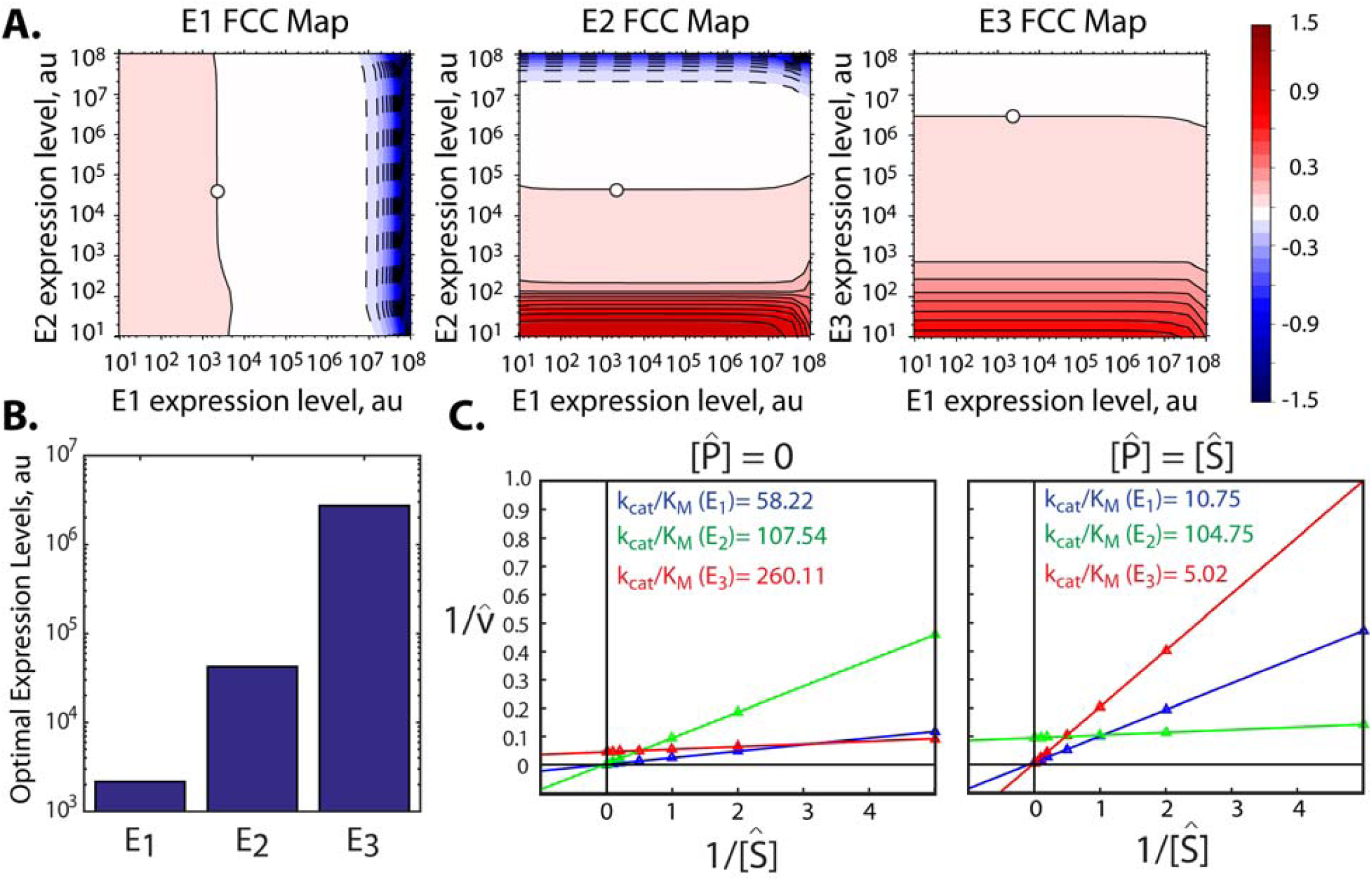
Pathway Mapping to Identify Rate-limiting Steps and Rank Enzyme Kinetics. (A) Flux control coefficient maps for a 3-enzyme pathway are shown. Stars indicate the optimal expression levels that are predicted to maximize the pathway’s overall flux. (B) Optimal enzyme expression levels as predicted by the Pathway Map Calculator. (C) Lineweaver-Burke analysis is used to determine the enzymes’ apparent k_cat_ and K_M_, using the kinetic metabolic model to simulate each enzyme’s reaction rate at varying substrate concentrations. Two scenarios are examined, where (left) the enzyme catalyzes its reactions by itself (out-of-pathway) or (right) the enzyme catalyzes its reactions within the overall pathway, reaching a non-equilibrium steady-state (in-pathway). The apparent k_cat_/K_M_ parameters in both scenarios are shown.(out-of-pathway) 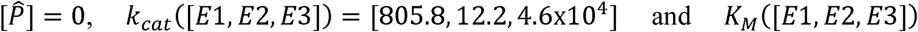 = [13.8,0.1,176.4]. and (in-pathway) 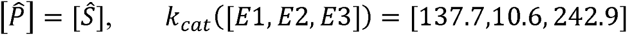 and 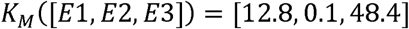.

However, when we applied the Pathway Map Calculator to identify the enzymes’ kinetic parameters (**Methods**), we found a noticeable difference in the apparent k_cat_ and K_M_ when the enzyme was treated as either out-of-pathway or in-pathway (**Figure 7C**). In the out-of-pathway scenario, measurements of the enzymes’ kinetic parameters are typically performed in *in vitro* reactions, starting in a reaction buffer that contains a selected amount of substrate, but without any product. When the analysis is performed using the out-of-pathway scenario, the Pathway Map Calculator finds that the enzyme E3 actually has the highest apparent k_cat_/K_M_ and E1 erroneously becomes the prioritized target for protein engineering. Instead, when the analysis is identically repeated using the in-pathway scenario, the Pathway Map Calculator correctly finds that the enzyme E3 has the lowest k_cat_/K_M_ of the three enzymes. The in-pathway scenario correctly takes into account the reversibilities of the reactions catalyzed by E2 and E3, which becomes important as product accumulates inside the cell. Notably, the algorithm identifies that E3 has a high apparent substrate K_M_ and a low apparent product K_M_, immediately suggesting that protein engineering efforts should modify the enzyme’s substrate binding pocket to decrease its substrate K_M_ and increase its product K_M_. Therefore, analysis using the in-pathway scenario should be used if protein engineering efforts are meant to improve the pathway’s overall productivity. Instead, if the apparent kinetic parameters are to be compared to standard *in vitro* reactions, then the analysis should use the out-of-pathway scenario. Importantly, this simple example reveals that measuring an enzyme’s k_cat_ and K_M_ in standard *in vitro* reaction conditions, in isolation of the pathway, can greatly mislead and misdirect protein engineering efforts when attempting to improve the overall pathway’s productivity.

## 3. Conclusions

The Pathway Map Calculator is a powerful computational algorithm with enormous potential to accelerate pathway optimization. We have demonstrated the algorithm’s capacity to accurately predict the expression-productivity relationship of many-enzyme pathways with limited and sparse experimental data, noisy end-product measurements, and containing allosteric regulation. The algorithm is broadly applicable to many industrially relevant pathways, and is agnostic to how the pathways are constructed and characterized, including the use of different types of genetic parts to vary enzyme expression, the expression of pathways in different organisms, and the measurement of metabolite productivities with different types of assays.

By regularly applying the algorithm to pathways of interest, we envision new modular mapping strategies whereby Pathway Maps of multiple pathways and networks are combined at the *model level* to predict more complex expression-productivity relationships, enabling precise tuning of expression to balance carbon, energy, and redox flows. Indeed, it is possible to combine Pathway Maps directly with the latest kinetic metabolic models of endogenous central metabolism (Chowdhury et al., 2014; Khodayari and Maranas, 2016; Khodayari et al., 2014) to predict how changes to endogenous enzyme expression levels will impact heterologous pathway productivities. When used in tandem with computational sequence design algorithms, such as the RBS Library Calculator and Operon Calculator (Farasat et al., 2014; Tian and Salis, 2015), the Pathway Map Calculator can predict the specific DNA sequences needed to maximize end-product biosynthesis. An online implementation of the Pathway Map Calculator can be found at https://salislab.net/software/PathwayMapCalculator.

## Acknowledgements

We would like to thank Zachary Costello, Hector Garcia Martin, and Jay Keasling for providing supplementary information on the limonene biosynthesis pathway as well as Alex Reis, Grace Vezeau, Chiam Yu Ng, and Amin Espah Borujeni for valuable discussion.

